# Protein intrinsically disordered regions have a non-random, modular architecture

**DOI:** 10.1101/2023.05.10.539862

**Authors:** Brendan S. McConnell, Matthew W. Parker

**Affiliations:** Department of Biophysics, University of Texas Southwestern Medical Center, Dallas, Texas 75235, USA

## Abstract

**Motivation:** Protein sequences can be broadly categorized into two classes: those which adopt stable secondary structure and fold into a domain (i.e., globular proteins), and those that do not. The sequences belonging to this latter class are conformationally heterogeneous and are described as being intrinsically disordered. Decades of investigation into the structure and function of globular proteins has resulted in a suite of computational tools that enable their sub-classification by domain type, an approach that has revolutionized how we understand and predict protein functionality. Conversely, it is unknown if sequences of disordered protein regions are subject to broadly generalizable organizational principles that would enable their sub-classification.

**Results:** Here we report the development of a statistical approach that quantifies linear variance in amino acid composition across a sequence. With multiple examples we provide evidence that intrinsically disordered regions are organized into statistically non-random modules of unique compositional bias. Modularity is observed for both low and high complexity sequences and, in some cases, we find that modules are organized in repetitive patterns. These data demonstrate that disordered sequences are non-randomly organized into modular architectures and motivate future experiments to comprehensively classify module types and to determine the degree to which modules constitute functionally separable units analogous to the domains of globular proteins.

**Availability and implementation:** The source code, documentation, and data to reproduce all figures is freely available at https://github.com/MWPlabUTSW/Chi-Score-Analysis.git. The analysis is also available as a Google Colab Notebook (https://colab.research.google.com/github/MWPlabUTSW/Chi-Score-Analysis/blob/main/ChiScore_Analysis.ipynb).

## 1 INTRODUCTION

The basic functional unit of a globular protein is the domain, a polypeptide region that folds into a stable three-dimensional structure (1). Many eukaryotic proteins possess a modular, multi-domain architecture resulting from the genetic duplication and shuffling (2) of the approximately 6,000 protein domain superfamilies (3). In this way evolution has produced a vast repertoire of architecturally distinct and functionally diverse multi-domain proteins. The domain architecture of a protein has traditionally been the realm of structural biology but modern bioinformatic algorithms now enable rapid identification of a protein’s separable domains (4). Understanding the modular architecture of multi-domain proteins has proven key to understanding their function.

Approximately 40% of the eukaryotic proteome does not fold into globular domains (5). These protein sequences, which do not possess stable secondary structure, exist in an ensemble of dynamic configurations and are described as being intrinsically disordered (6). Despite their lack of structure, protein intrinsically disordered regions (IDRs) are known to play essential roles in many physiological and pathophysiological pathways, including transcription (7,8), DNA replication (9), as the etiological agents in certain neurodegenerative proteinopathies (10), and certain IDRs drive protein phase separation which helps organize the cell (11). Regions of protein disorder can be discriminated from globular domains by sequence composition alone (12,13) and there exist many bioinformatic algorithms to predict the disorder propensity of a polypeptide (14).

By definition, IDRs lack a defined spatial architecture and, with the exception of short linear motifs (SLiMs) (15), are thus relatively unrestrained in primary structure. As a result, there are generally lower levels of conservation between orthologous IDRs compared to folded regions (16,17) and IDR primary structure often appears to lack organization. One exception to this are seemingly non-random regions of disordered sequences that are locally enriched in only a small subset of amino acids, so-called Low Complexity Regions (LCRs). Sequence gazing readily identifies LCRs by their conspicuous local sequence bias but these can also be quantitatively and unbiasedly discriminated on the basis of informational entropy (18). Some IDRs also possess multiple LCR types with distinct functionalities (19,20) and bioinformatic approaches to demarcate unique LCR subsequences on the basis of their composition and amino acid dispersion have recently been reported (20,21). LCRs, however, represent but a fraction of all disordered sequences and it is unknown if sequence-spanning organizational principles are operative in IDRs generally.

The sequence bias observed in many IDR sequences has motivated the development of bioinformatic algorithms to classify disordered protein sequences on the basis of composition. These approaches have focused on annotating IDRs and their sub-regions according to a limited set of *known* compositional varieties (i.e., “flavors”), such as by charged residue content (e.g., polyampholyte or strong polyelectrolyte) (22,23) or bias for a given amino acid type (e.g., polar residues) (24,25). Guided by this concept, sequence analysis tools have been built that will demarcate regions within a protein that match a user-defined composition (26). Although useful, these techniques require *a posteriori* knowledge of IDR flavors. In this sense, these methods are *candidate-based* analysis tools, being very good at determining whether a sequence is or isn’t of the candidate class but not useful in identifying new compositional varieties that exist undiscovered within a sequence.

Here we report the development of a statistically robust computational algorithm that unbiasedly maps compositionally-distinct subsequences within an IDR. Relying on the Chi-Square (χ^2^) Test of Homogeneity, our Chi-Score Analysis quantifies variability in the fractional composition of amino acids between two sequences. Applied intramolecularly in a moving-window, matrix-based approach, this method can identify sequence-spanning compositional heterogeneity to parse a protein sequence into regions of distinct amino acid composition, regardless of what those compositions are. With multiple examples we show that IDRs of both low and high sequence complexity possess local compositional bias that bestows disordered sequences with a non-random, modular architecture. Analogous to the domain architecture of globular proteins, we propose that modules (i.e., compositionally distinct subsequences) represent functionally separable units of disordered sequences, and our unbiased, discovery-based approach to their identification represents a promising new direction in IDR classification. Altogether, these data demonstrate that high-level organizational principles are at work in disordered sequences and motivate future functional studies to understand the role of modules in biology.

## 2 RESULTS & DISCUSSION

### 2.1 Development of a bioinformatic algorithm to identify local compositional bias in IDRs

Protein disordered regions lack stable secondary structure and are thus relatively unrestrained in primary structure. Consistently, IDRs often have weak linear sequence conservation (17,27) and no visually discernible sequence patterns, suggesting they lack a defined organization. We hypothesized that local compositional bias may bestow disordered sequences with a sequence-spanning level of organization undetectable by current methods. We therefore developed a bioinformatic algorithm to determine in an unbiased and quantitative fashion if amino acids are non-randomly distributed across the length of a protein sequence and to map this information back onto the sequence. Our approach implements the Chi-Square (χ^2^) test statistic to compare the fractional content of amino acids between two sequences as a measure of compositional dispersion (see **METHODS AND ALGORITHMS**). Applied in this way, the chi-score quantifies how different the amino acid proportions (i.e., composition) are between two sequences.

The chi-score metric can be applied intramolecularly to identify compositionally biased regions (**Figure 1A**). In the first step of our algorithm, the sequence is broken into all possible subsequences for a specified window size and all pairwise chi-scores are calculated (**Figure 1A-a**). Each chi-score is then converted to a Pearson’s correlation coefficient which better resolves compositionally biased regions and sequence-spanning patterns (**Figure 1A-b**). Subsequently, the mean correlation coefficient of each subsequence is calculated from a subsequence-centered square window and these “insulation scores” are plotted against residue position (**Figure 1A-c**). Finally, local minima from the insulation score plot are calculated and recorded as potential boundaries between compositionally distinct regions (**Figure 1A-d**). These steps are completed for nine sets of pairwise chi-scores – each using a different even integer window size between 6 and 22 to define the original subsequences – which results in nine sets of boundaries (**Figure 1A-d**).

**Figure 1:**
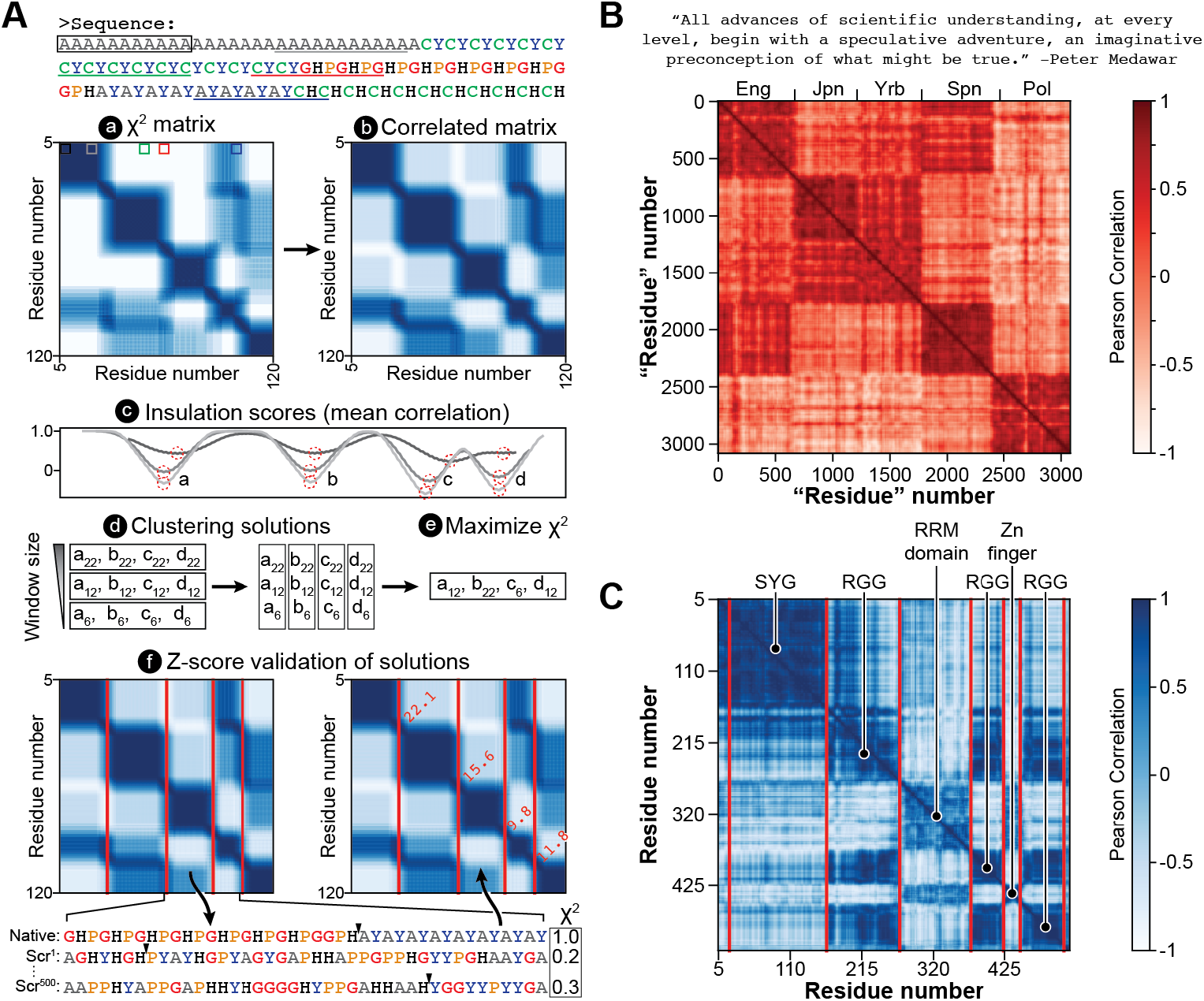
A bioinformatic algorithm to measure compositional dispersion across a sequence. A) For a given sequence, a) pairwise chi-scores are calculated for all window-defined subsequences and plotted as a matrix. Window sizes include every even size from 6-22. Then, for each window size, b) chi-scores are converted to Pearson’s correlation coefficients, c) insulation scores are calculated to define local minima which represent compositional boundaries, and d) the minima identified for all window sizes are grouped according to proximity. From these groups, e) each boundary placement is optimized to maximize chi-score and f) z-scores are calculated for each boundary element and are used to iteratively remove non-significant boundaries. B) The Chi-Score Analysis method can distinguish between human languages on the basis of alphabet usage. A quote from Peter Medawar’s Romane Lecture was translated into five languages and each translation was strung end-to-end for analysis. “Eng” = English, “Jpn” = Japanese, “Yrb” = Yoruba, “Spn” = Spanish, and “Pol” = Polish. C) The Chi-Score Analysis method can distinguish between regions of sequence bias in the human protein FUS. Boundaries are shown for 95% confidence level.

The boundaries, which were originally grouped by window size, are then clustered by residue proximity (**Figure 1A-d**) and the optimal boundary positions are determined by selecting the placements within each group that maximize the mean chi-score between the surrounding regions (**Figure 1A-e**). The statistical strength of each boundary is then determined by calculating a z-score. To do this, the two juxtaposed regions separated by each boundary are scrambled 500 times and the boundary position resulting in the maximum chi-score identified. This results in a set of 500 chi-scores, one for each scramble, from which the mean (and standard deviation) is determined and used to calculate the z-score for the corresponding boundary (**Figure 1A-f**). Finally, low scoring boundaries are iteratively removed and those that remain are re-optimized and scored until only high-confidence boundaries remain.

To demonstrate the utility of this algorithm in identifying compositional bias, we first tested its effectiveness at differentiating human languages (**Figure 1B**). We translated a quote into English (Eng), Japanese (Jpn), Yoruba (Yrb), Spanish (Spn), and Polish (Pol), appended the translations one after another, and then analyzed the resulting string of text with the Chi-Score Analysis. This approach proved highly effective at discriminating between languages based on their differential character usage (all texts were Romanized and composed of the same 26 letter alphabet).

Regions off the diagonal with relatively high correlation reveal languages with more similar alphabet usage, such as English and Spanish or Yoruba and Japanese. Conversely, regions off the diagonal with relatively low correlation reveal languages with differential alphabet usage, such as Polish and Yoruba. We next applied the method to a protein sequence to determine whether it can parse a sequence by “molecular language” (**Figure 1C**). Fused in Sarcoma (FUS) is known to possess multiple compositionally biased regions with unique functionality (28,29) and our method accurately identifies these, revealing three major language types: G/S-Y-G/S repeats, RGG repeats, and a more complex sequence type which corresponds to the folded domains of the protein (RRM and Zn finger domains). Analysis of off-diagonal correlated regions reveals homology amongst the three RGG-enriched regions and the two regions which have a globular structure. These data establish the utility of the Chi-Score Analysis in parsing sequences by local compositional bias.

### 2.2 IDRs have a non-random, modular organization

Having established the algorithm on a sequence with conspicuous local sequence bias (**Figure 1C**), we next applied it to a disordered region that lacks recognizable sequence patterning. The Origin Recognition Complex (ORC, composed of Orc1-6) is an essential DNA replication initiation factor that contains an IDR in the Orc1 subunit that is necessary for recruitment to chromatin (9). In *C. elegans*, the Orc1 IDR is predicted to be 245 amino acids long and, except for two short, low complexity regions, has no apparent sequence organization (**Figure 2A**). However, Chi-Score Analysis reveals a strikingly non-random, sequence-spanning level of organization with regions of distinct compositional bias juxtaposed in a repetitive pattern (**Figure 2B**). Given the modular appearance, we will refer to these compositionally biased regions as “modules”. The Orc1 IDR has module types which can be loosely categorized as either basic, neutral, or acidic. In this sequence the basic modules (residues 1-51, 106-167, 186-232) and acidic modules (residues 52-68, 80-105, 168-185, 233-245) alternately repeat throughout the sequence, with the neutral module type appearing only once (residues 69-79). While charge-based classification is convenient, a more careful investigation of module sequences (**Table 1**) reveals greater complexity than a simple alternation of charge, with non-charged residues also being differentially patterned. Notably, the majority of Orc1 IDR modules identified by the Chi-Score Analysis are not annotated in Uniprot as having “Compositional bias”, emphasizing the importance of our discovery-based approach (see **Supplemental File 1** for a list of module sequences and a comparison with Uniprot annotations).

**Figure 2:**
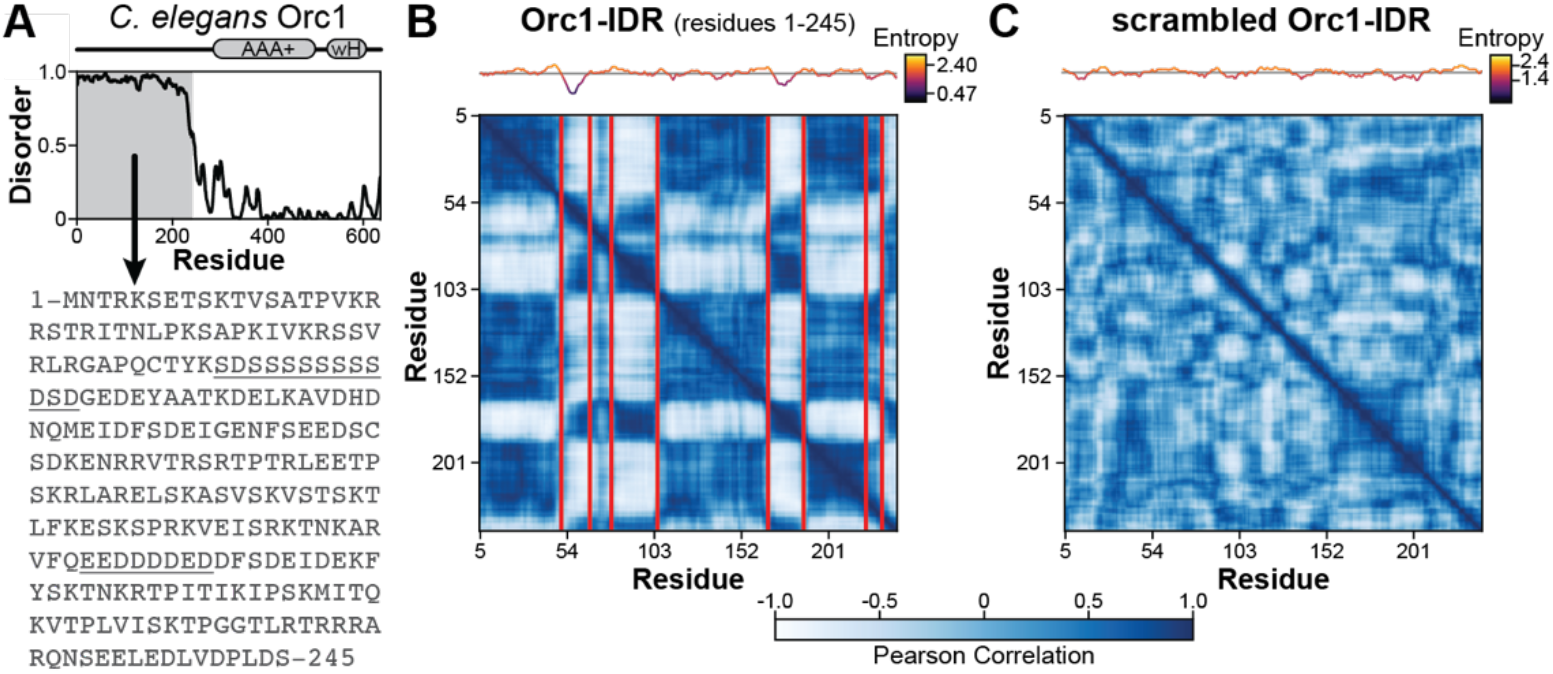
High-complexity disordered regions are non-randomly organized. A) The *C. elegans* Orc1 IDR (Orc1-IDR) has a long N-terminal IDR (residues 1-245) as predicted by Metapredict (43) but no obvious sequence patterning. B) Chi-Score Analysis reveals a strikingly modular architecture for Orc1-IDR with alternately repeating blocks of like-type sequences. Boundaries are shown for 95% confidence level and a sequence entropy plot is shown above the matrix. C) Modularity is lost when the sequence of Orc1-IDR is randomly scrambled.

**Table 1:**
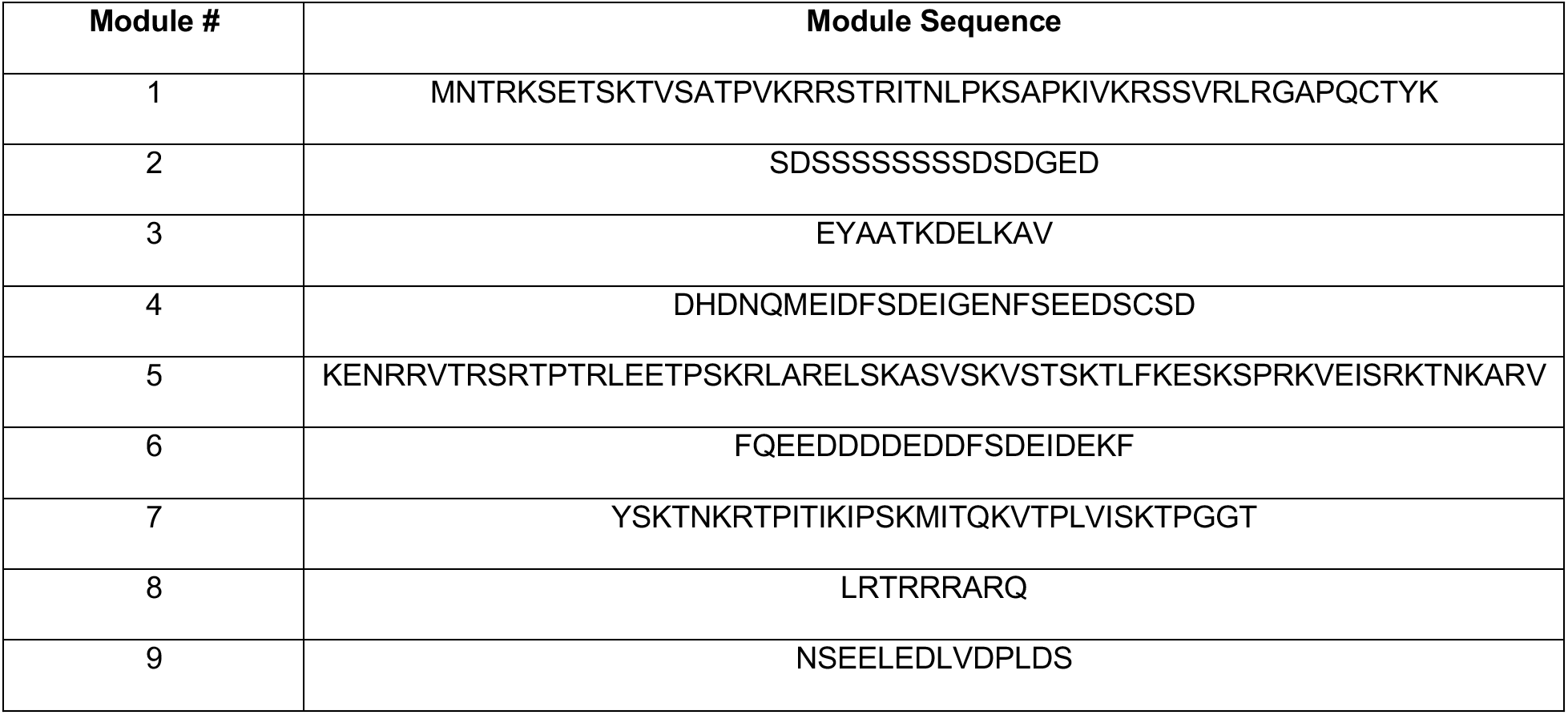
Module sequences for the Orc1 IDR.

The discovery of a statistically non-random, sequence-spanning level of organization in the Orc1 IDR was unexpected. Importantly, amino acid content alone does not produce this type of organization. To demonstrate this, we reran the analysis on a random scramble of the Orc1 IDR (**Figure 2C**). In this example, modules are not only visually absent but there were additionally no statistically significant boundaries output by our algorithm. Likewise, compositionally biased modules are absent from Orc1’s folded domains (**Figure 2 – figure supplement 1A**) which, compared to the IDR, have a far more uniform sequence landscape. This is consistent with prior work showing that the lengthwise distribution of amino acids in globular domains does not differ substantially from randomized sequences containing equivalent proportions of amino acids (30–32). These data suggest that local compositional bias is an organizational principle unique to disordered sequences.

**Figure 2 – figure supplement 1:**
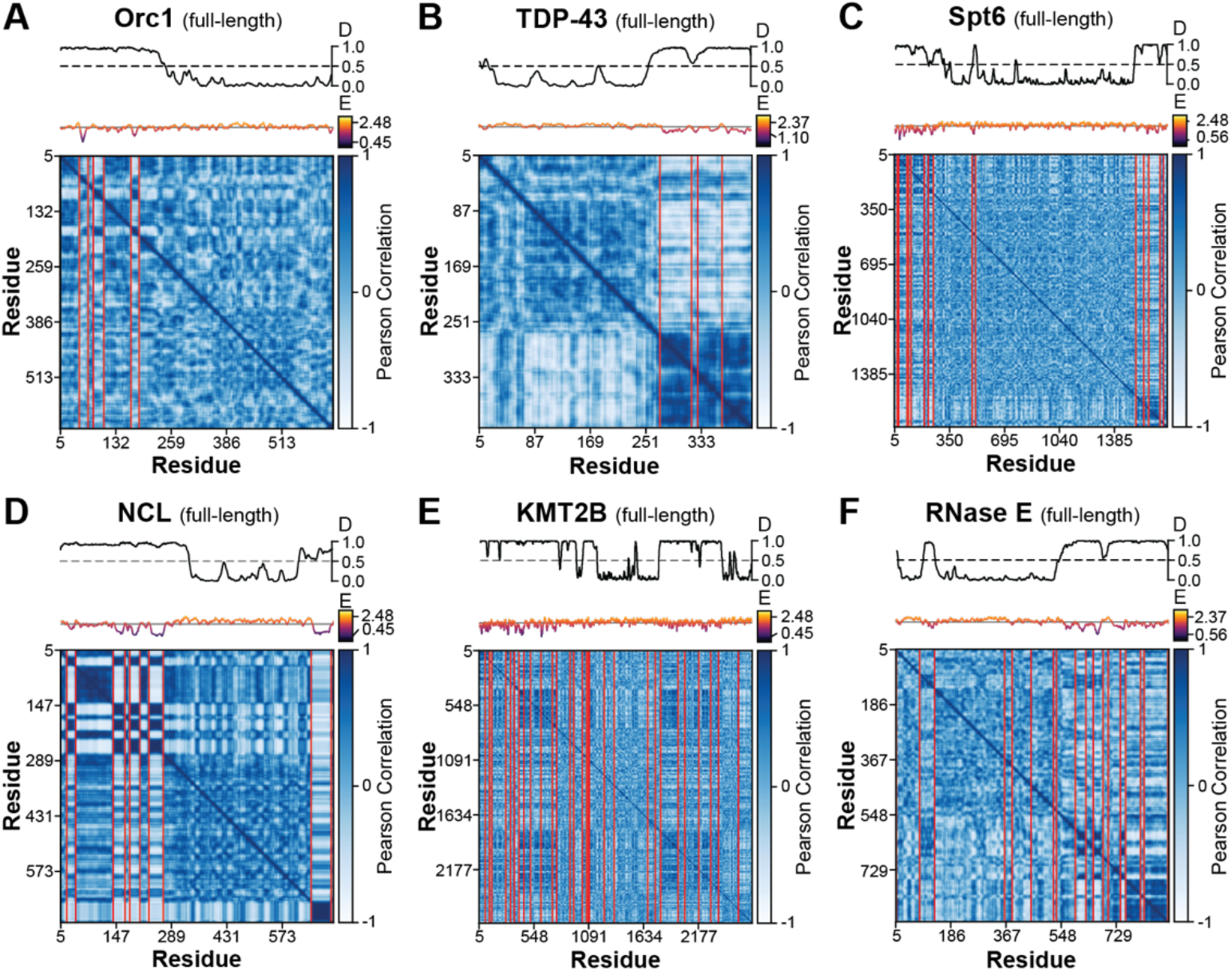
Compositionally-defined modules are largely absent from protein folded domains. Chi-score analysis for full-length A) *C. elegans* Orc1, B) human TDP-43, C) human Spt6, D) human Nucleolin (NCL), E) human KMT2B, and F) *C. crescentus* RNaseE. Boundaries are shown for 95% confidence level and a sequence disorder (“D”) and entropy (“E”) plot are shown above each matrix.

### 2.3 Many disordered sequences have local compositional bias

The strikingly modular architecture of FUS (**Figure 1C**) – a low complexity IDR – as well as Orc1 (**Figure 2B**) – a high complexity sequence – prompted us to investigate whether this type of organization is widely operative in disordered sequences. We therefore extended our studies to assess modularity of several other protein disordered regions with known biological and pathological significance, including human TDP-43 (**Figure 3A**), Spt6 (**Figure 3B**), Nucleolin (NCL, **Figure 3C**), KMT2B (**Figure 3D**), and *Caulobacter crescentus* Ribonuclease (RNase) E (**Figure 3E**). These analyses, which we briefly describe below, show that local compositional bias is pervasive amongst IDRs and bestows disordered sequences with a modular architecture. Conversely, local compositional bias appears largely absent from folded domains, at least for the proteins under consideration here (analysis of full-length sequences is shown in **Figure 2 – figure supplement 1B-F**).

**Figure 3:**
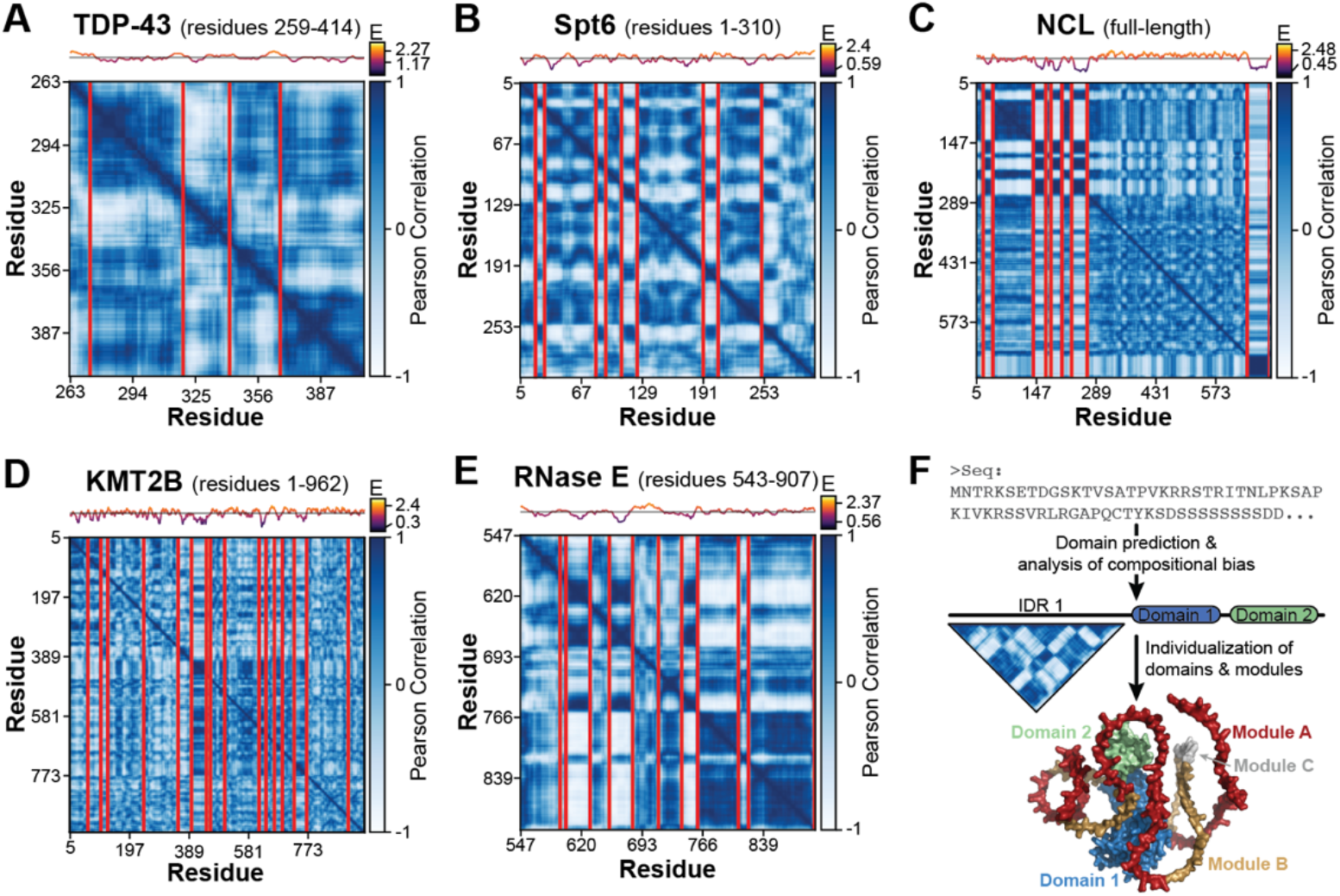
Many intrinsically disordered regions have a modular architecture. Chi-Score Analysis for A) human TDP-43 IDR (residues 259-414), B) human Spt6 N-IDR (residues 1-310), C) human Nucleolin (NCL), D) human KMT2B N-IDR (residues 1-962), and E) *C. crescentus* RNaseE C-IDR (residues 543-907). Boundaries are shown for 95% confidence level and a sequence entropy (“E”) plot is shown above each matrix. NCL contains four tandem RNA binding domains which are contained within the largest module in the matrix and appear compositionally uniform. F) IDRs should not be thought of as indivisible units but as modular sequences with region-specific physicochemical features and biological function.

We analyzed an internal IDR of the protein TDP-43 (residues 259-414, **Figure 3A**), an RNA processing factor that forms pathological aggregates in neurodegenerative disease. Our analysis identified five modules, one of which corresponds precisely to a region harboring an abundance of pathological missense mutations and which functions independently as TDP-43’s amyloidogenic core (residues 320-PAMMAAAQAALQSSWGMMGMLAS-342) (33–36). These data suggest that modules, like a folded domain, can behave as functionally separable units. Interestingly, none of the TDP-43 modules identified by our analysis are annotated in Uniprot (**Supplemental File 1**). Chi-Score Analysis of the N-terminal IDR of Spt6 reveals several highly distinct modules (residues 1-310, **Figure 3B**) which, together with the prior literature (8), suggests that modules can also demonstrate emergent behavior, with their functionality derived from the cooperative interactions of modules. Specifically, we identified a repetitive pattern of basic and acidic modules, and recent data show that these alternating blocks of charge mediate selective partitioning of Spt6 into MED1 condensates to control transcriptional activation (8). We anticipate that understanding module types and patterning will help elucidate the rules underlying selective partitioning in biomolecular condensates. This idea is supported by Chi-Score Analysis of full-length Nucleolin (NCL, **Figure 3C**), where we identify a pattern of modules within its N-terminal IDR that was recently shown to mediate the protein’s selective partitioning into the nucleolus (37). Finally, we identified numerous modules of variable composition and size within the N-terminal IDR of KMT2B (**Figure 3D**) and the C-terminal IDR of RNase E (**Figure 3E**). Module sequences for each protein can be found in **Supplemental File 1**.

Altogether, these data suggest that local compositional bias represents an organizational principle that is widely operative in disordered sequences and inspires fundamental questions about the role of IDR modularity in biology and disease. A significant advantage of the Chi-Score Analysis method is that compositionally-distinct regions can be discovered solely on the basis of amino acid bias without user-defined search criteria. Even in the small set of sequences analyzed here, this unbiased approach to module identification suggests a level of IDR compositional diversity (i.e., flavors) that dwarfs existing classification paradigms (**Supplemental File 1**).

## 3 CONCLUSION

We find that the sequences of both low and high complexity IDRs are non-randomly organized into regions with local compositional bias. This concept, like the structural hierarchy (1° – 4° structure), bears the hallmarks of a fundamental organizational principle: it is generalizable to any sequence, it appears to be broadly operative, and it enables the unbiased sub-division of a sequence into component parts. These features have enabled the structural hierarchy to provide a comprehensive classification of folded sequence space, a long-standing objective in the field of protein disorder. Collectively, these findings warrant a conceptual shift away from IDRs as indivisible functional units to viewing them as modular sequences with region-specific physical, chemical, and functional properties (**Figure 3F**). To this extent, individual IDR modules and their combinations may represent the functional analog of the globular protein’s domain, whose many types and arrangements produce the diversity of cellular functions that are needed for life. This concept has empirical support from studies of transcription factor transactivation domains (7) and prion domains (37,38) where sub-sequences in extended regions of disorder contribute specific functionality strictly on the basis of their composition.

This work calls for the comprehensive classification of module types and their combinations to determine whether there exist distinct classes or a continuum of compositional varieties. Such work will benefit from other sequence characterization parameters, such as charge distribution (22) and binary sequence patterning (39). While the evolutionary preservation of modules provides strong *a priori* evidence that modules are important for biology, future studies are needed to systematically relate module type with functionality. Beyond this, many other important questions remain, including the use of genetic mechanisms to shuffle modules and produce novel functionalities, the biophysical properties of isolated modules and emergent properties of multi-module sequences, and whether a modular view of IDRs clarifies the mechanism of disease-associated mutations and rationalizes specificity in IDR-enriched biomolecular condensates. In the immediate, we hope the concept of IDR modularity as a generalizable organizational principle serves as a framework for generating hypotheses for IDR functional mechanisms and provides a rational approach for dissecting this enigmatic class of sequences (**Figure 3F**). To facilitate these types of studies, the code required to run these analyses is freely available and easily implemented (see **METHODS AND ALGORITHMS**).

## 4 METHODS AND ALGORITHMS

### 4.1 Applying the Chi-square test to compare sequence composition

The Chi-Square Test of Homogeneity is used to determine whether two distributions are from, or were sampled from, the same population. Traditionally, the Chi-Square test statistic is calculated and used to either reject or accept the null hypothesis. Here, the test statistic is instead used as a metric scoring the compositional difference between two sequences; a high-test statistic indicates a high degree of compositional distinction.

The Chi-square test can be applied both intermolecularly, comparing the amino acid content of different protein sequences, and intramolecularly, comparing the amino acid content of subsequences within a single protein. This latter application, which is elaborated on in the following section, applies a matrix-based approach to parse a sequence into compositionally distinct regions from pairwise subsequence comparisons. When two sequences are compared, the number of each residue is first taken as the observed values (O) for the Chi-Square formula. For each observed value, a corresponding expected value (E) is calculated with the following equation.

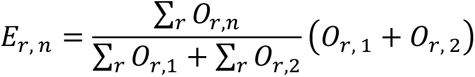

In this formula, *n* refers to the sequence (either 1 or 2) and *r* refers to the residue (one of twenty amino acids). To determine the expected value for alanine residues in the first sequence, the total number of alanine residues in the two sequences is multiplied by the ratio of that sequence’s length to the total length of both sequences. The test statistic can then be calculated with each observed/expected pair using the following equation.

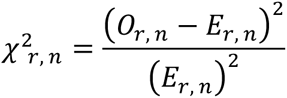

Finally, the 40 unique scores – one for each amino acid and sequence pair – are summed, and the test statistic is normalized between zero and 1. This is done by dividing this sum by the maximum possible score for those two sequences, which is equal to the sum of their lengths and occurs when they have no residues in common. Sequences that have no residues in common will always receive a normalized score of one, and sequences with identical amino acid compositions will receive a normalized score of zero.

In addition to the chi-score method, Euclidean distance has also been applied to quantitate the similarity in amino acid composition between disordered sequences (40,41). This method takes the fractional content of amino acids as Cartesian coordinates in a 20-dimensional Euclidean space and the “distance” between any two sequences is quantified. Our chi-score method builds on Euclidean distance in several ways. First, chi-score values can be readily decomposed to see the contribution of each amino acid to the overall score, thereby determining the residue type(s) that most distinguishes one sequence from another. Second, Euclidean distance is influenced by sequence complexity while chi-score is not. Finally, the Chi-Square test possesses inherent statistical power which we apply to determine whether subsequences are compositionally distinct compared to random scrambles of the same sequence.

### 4.2 Applying the Chi-Score Analysis intramolecularly to identify regions of local compositional bias

The Chi-Score Method can be applied intramolecularly to identify regions of distinct amino acid compositional bias. During the first step of this analysis, the sequence is broken up into all possible subsequences of nine different window sizes (all even integers between 6 and 22) and for each window size the chi-score is calculated for all subsequence pairs. The pairwise scores are then converted to Pearson’s Correlation Coefficients with the following equation:

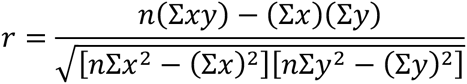

After plotting the coefficients as a two-dimensional matrix with the subsequence positions along each axis, we then calculate insulation scores (as used in Hi-C data processing (42)) for each residue position. These scores are calculated by taking the average value in a square window that slides along the main diagonal. Insulation score values are high when this square is within a region of distinct amino acid composition and low when the square is near the boundaries between them. Therefore, we then take the residue positions of the local minima of these insulation scores as potential boundaries between compositionally distinct modules.

Because this is done with nine different window sizes, we now have multiple sets of potential boundaries grouped by the window size used to calculate them. Modules of different length and/or complexity will be best identified with different window sizes, so the precise positions for each boundary will vary between the nine groups. We then regroup these boundaries spatially so that they are clustered with other potential positions for the same boundary. The optimal position for each boundary is then determined by selecting those that maximize the chi-scores between the resulting modules.

The boundaries can now be statistically verified by calculating a z-score for each. First, the two modules separated by a boundary are juxtaposed and randomly scrambled 500 times. Then, the greatest chi-score between two modules that can be achieved with a comparable boundary is determined for each scramble. Finally, these scores are used to convert the raw chi-score for that boundary into a z-score, which tells us how likely it is that the predicted boundary separates truly distinct modules or simply identifies local biases occurring by chance. Boundaries with low z-scores are iteratively removed until only those with significant scores remain. After a boundary is removed, the positions are reoptimized and the z-scores are recalculated for those that remain; the placements and z-scores corresponding to each iteration are also stored so that they can be recalled as desired.

### 4.3 Python implementation and accessibility

The code to perform these analyses was designed with Python Version 3.10.8 and executed in Jupyter Notebook. To allow for easy access and implementation of the analysis, we have made it freely available in a number of formats: 1) the source code, which contains all functions necessary to perform and manipulate the analysis as desired, 2) a streamlined Python notebook that installs the algorithm and performs the analysis on a single protein sequence, 3) a Python notebook that performs the analysis on a step-by-step basis so that the outputs of each can be recalled as desired, and 4) a Google Colab notebook that lets the user input a sequence and adjust optional parameters. For the easiest implementation of this analysis, we recommend using the Google Colab notebook, which can be found at: https://colab.research.google.com/github/MWPlabUTSW/Chi-Score-Analysis/blob/main/ChiScore_Analysis.ipynb. For instructions on how to use the other implementations, as well as the code required to reproduce all matrices shown throughout the paper, please see: https://github.com/MWPlabUTSW/Chi-Score-Analysis.git.

## ACKNOWLEDGEMENTS

We thank Xiaochen Bai, Nick Grishin, Michael Rosen, and Weiwei Wang for critical reading of the manuscript. M.W.P. is the Cecil H. and Ida Green Endowed Scholar in Biomedical Computational Science. This work was supported by The Welch Foundation (I-2074-20210327, to M.W.P.), the Cancer Prevention and Research Institute of Texas (CPRIT, RR200070, to M.W.P.), and the National Science Foundation (NSF, 2308642, to M.W.P). The funders had no role in study design, data collection and analysis, decision to publish, or preparation of manuscript.

